# How reliable is the link between effort and nucleus accumbens connectivity at rest? A psychometric evaluation of repeated assessments

**DOI:** 10.64898/2026.11.19.767858

**Authors:** Zsófia Karlócai, Anne Kühnel, Corinna Schulz, Melina Grahlow, Kristin Kaduk, Andreas Frick, Birgit Derntl, Nils B. Kroemer

**Author notes:** **Corresponding author:** Prof. Dr. Nils B. Kroemer, Venusberg Campus 1, 53127 Bonn, Germany.

## Abstract

Individuals differ in their willingness to work for rewards. Although effort-based behavior shows trait-like stability and motivational traits associate with functional connectivity (FC), it remains unclear how robustly individual differences in effort are reflected in intrinsic FC of reward circuits. Here, we investigated whether effort-based behavior and questionnaire scores are reflected in nucleus accumbens (NAcc) FC. To this end, we evaluated this association in 84 healthy adults (43 women, 41 men), who completed five sessions of an effort allocation task and two task-free fMRI scans, each collected on different days to capture trait-like behavior and brain function. First, behavioral measures showed high test-retest reliability across sessions (ICCs: 0.62– 0.83). Second, NAcc FC showed the highest reliability in subcortical regions (ICCs = 0.59–0.69), while cortical connections were heterogeneous and, overall, less reliable. Higher effort exertion was associated with stronger NAcc<>IFG FC (*t*_max_(76) = 4.65, *p*_FWE_ = .041, *k* = 141), while greater sensitivity to task difficulty was associated with stronger NAcc<>subcallosal gyrus FC (*t*_max_(76) = -5.15, *p*_FWE_ = .028, *k* = 155). Intriguingly, the association of effort exertion with NAcc<>IFG FC decreased over behavioral sessions, whereas the association of sensitivity to cost (difficulty) with NAcc<>subcallosal FC was stable over behavioral sessions. Our findings show that identifying robust brain–behavior associations likely require comparable multi-session approaches, at least in moderately sized samples. Beyond reliability, such approaches may also help reveal additional insights about the temporal profile of brain–behavior associations, which may differ markedly for new versus routine task performance.

## Introduction

Every day, we face choices that weigh effort against potential rewards, whether to go for a run after a long day or rest on the couch. This willingness to invest effort to obtain future rewards is a fundamental aspect of human behavior (Rigoli & Pezzulo, 2022). Effort-based decision-making tasks formalize this as the integration of expected effort costs and benefits to guide goal-directed action (Treadway et al., 2012). Disruptions in this integration are thought to underlie motivational impairments, such as apathy and anhedonia, which are observed across a wide range of mental and neurological disorders, including depression and schizophrenia (Chang et al., 2019; Husain & Roiser, 2018; Randez & Hélie, 2024), and are associated with worse clinical outcomes (Van Reekum et al., 2005; Vinckier et al., 2017). Even healthy individuals vary markedly in their willingness to exert effort (Chong et al., 2016; Reddy et al., 2015), and this variation has been linked to differences in daily productivity and initiative (Bonnelle et al., 2015; Hewitt et al., 2025) as well as striatal dopamine function (Treadway et al., 2012; Westbrook et al., 2020). Individual differences in effort exertion may partly reflect variation in brain networks involved in evaluating rewards and effort costs and translating these evaluations into motivated action (Trinh et al., 2026). In laboratory settings, motivational decision-making is often measured with effort tasks, such as discounting or allocation paradigms. In such tasks, participants choose between options that vary in reward magnitude and required effort (Klein-Flügge et al., 2016; Salamone et al., 2015; Westbrook et al., 2013). Behavior is often highly test-retest reliable in individuals, rather than contextual (Malik et al., 2025; Ohmann et al., 2022; Van Den Hoek Ostende et al., 2021). Accordingly, individual differences in effort-based decision-making may provide a useful phenotype for studying trait-like differences in motivation, including associations with brain function within motivational circuits. Such trait-like differences in brain function can be captured by functional connectivity (FC) profiles, which are so highly reliable to allow re-identification of individuals across independent measurements (Finn et al., 2015). Consequently, they have been linked to motivational traits such as reward sensitivity, impulsivity, and apathy in healthy adults (Adrián-Ventura et al., 2019; Derosiere et al., 2025; Xin et al., 2020; Zhao et al., 2025). However, whether seed-based FC reliably tracks trait-like individual differences in effort-based behavior remains to be established.

Reliability is a prerequisite for diagnostic measures and test-retest assessments are crucial to determine stability (Aldridge et al., 2017; Fröhner et al., 2019; Hedge et al., 2018). Test-retest reliability can be quantified with the intraclass correlation coefficient (ICC), with values below 0.4 considered poor and above 0.6 good (Shrout & Fleiss, 1979). Measures with low ICCs cannot reliably discern individual differences, limiting their utility for brain–behavior association research. Effort-discounting measures show moderate to good stability over repeated testing (Escobar et al., 2023), and this pattern extends to effort tasks more broadly. The high test-retest reliability of effort tasks even holds across naturalistic settings, where additional state-like variability has been shown (Hewitt et al., 2025). In contrast, lower overall ICCs have been observed for computational parameters of reinforcement learning (Kühnel et al., 2025; Mkrtchian et al., 2023; Neuser et al., 2023). Self-reported measures of motivational traits, such as apathy, anhedonia, and reward sensitivity, also capture trait-like individual differences in motivation with high reliability (Husain & Roiser, 2018; Lockwood et al., 2017; Treadway et al., 2009), but they are only weakly associated with task-based measures of motivation (Renz et al., 2023). Whether effort-based tasks and motivation questionnaires correspond distinctively to FC of the motivational circuit therefore remains an open question.

Neuroimaging studies have highlighted the role of cortico-striatal circuits in motivated behavior, particularly connections between the striatum and prefrontal areas such as the ventromedial prefrontal cortex (vmPFC) and anterior cingulate cortex (ACC) (Ballard et al., 2011; Holroyd & Yeung, 2012; Li et al., 2018; Siemsen et al., 2025), which integrate reward value, effort cost, and anticipated outcomes during goal pursuit. Resting-state FC (rsFC) within reward-related circuits has been linked to individual differences in motivation. For example, higher reward sensitivity was associated with stronger connectivity between the ventral tegmental area and vmPFC, as well as between the vmPFC and ACC, in healthy adults(Adrián-Ventura et al., 2019). Altered FC in these circuits has also been linked to reward motivation adaptation in individuals with negative schizotypal traits (i.e., subclinical characteristics involving reduced pleasure, motivation, and social engagement; Yan et al., 2024), and impulsivity-related alterations in substance use disorder (Zhang et al., 2025). Within this circuitry, the nucleus accumbens (NAcc) is of particular interest because of its central role in reward processing, goal-directed action, and effort-based motivation (Salamone & Correa, 2024; Yan et al., 2023). NAcc FC has also been linked to individual differences in reward-related behavior in healthy adults, with stronger NAcc<>orbitofrontal rsFC associated with better reinforcement learning (Kaiser et al., 2018), and FC during a monetary incentive delay task was associated with reward sensitivity and behavioral motivation (Wyngaarden et al., 2026). In a large clinical sample, NAcc FC profiles have been associated with symptom-specific differences in Δappetite, reflecting increases or decreases in appetite and weight in patients with depression (Kroemer et al., 2022), illustrating that NAcc FC captures clinically relevant differences in appetitive behavior. Such studies often rely on a single fMRI session, although whole-brain rsFC profiles often only show moderate test-retest reliability, particularly if striatal seeds are used (Noble et al., 2019). Consequently, large-scale brain-wide association studies indicate that associations between inter-individual differences in brain measures and behavior are often small and sensitive to sampling variability (Marek et al., 2022). Thus, although reliable behavioral and neural measures are a prerequisite for robust brain–behavior associations, it is not clear whether repeated sampling can help overcome the limited power afforded by moderate sample sizes.

To address this gap, we combined repeated behavioral assessment across five sessions with two task-free fMRI measurements each collected on different days. Our goal was to characterize the test-retest reliability of effort-based behavior and NAcc FC to then test whether reliable individual differences in effort indices are associated with NAcc FC patterns. Then, we assessed whether these NAcc-behavior associations are specific to task-based behavioral measures compared to questionnaire-based measures of motivational traits. Our findings indicate that effort indices and NAcc FC show sufficient test-retest reliability to capture robust individual differences. Building on this insight, we observed individual differences in the general tendency to exert effort associated with NAcc<>IFG FC and effort cost sensitivity associated with NAcc<>subcallosal gyrus FC. Notably, these associations were most pronounced for effort-based behavior and did not emerge for subjective wanting ratings or questionnaires. This highlights that NAcc FC profiles can be used as a proxy of trait-level differences in individual motivation to exert effort.

## Methods

The reported data are part of a larger study examining the association between metabolic states and reward learning, preregistered on the Open Science Framework (https://osf.io/gpr52). The specific analyses were not preregistered, but were designed to evaluate the psychometric characteristics for planned analyses. The study was previously described in Kaduk et al. (2026) and in Grahlow et al. (2026).

### Participants

We initially included 97 participants in the study. Of these, 93 completed all five behavioral sessions and were included in the behavioral reliability analyses. Nine participants dropped out after the behavioral assessments and had to be excluded due to missing neuroimaging data, resulting in a total of 84 participants for the analyses (43 females, 41 males, whole sample: *M*_age_ = 24.04 ± 3.44, range: 18-32 years; *M*_BMI_: 24.89 ± 4.01 kg/m^2^, range: 18-36.7 kg/m^2^). The primary brain–behavior analysis included 84 participants, whereas the replicability analyses of brain–behavior associations excluded one participant because of incomplete fMRI data (only one session). Apart from an elevated BMI range, participants were physically and mentally healthy (according to self-reports). Eligibility was determined via telephone screening. Exclusion criteria included self-reported BMI <18.5 or >35, serious somatic illnesses, and current or past-diagnosed mental disorders such as bipolar disorder or schizophrenia. Depression, eating disorders of any kind, anxiety or panic disorders, obsessive-compulsive disorder, and somatization disorders must have occurred >12 months ago. As the study also investigated the effects of fluctuations of sex hormones across the menstrual cycle, women currently using hormonal contraception were also excluded. Likewise, pregnancy, breastfeeding, irregular hormonal cycles, or premenstrual dysphoric disorder were exclusion criteria. MRI compatibility was confirmed for all participants. Participants provided written informed consent and received performance-based monetary and food rewards plus a fixed compensation of €160 for completing the study. The study was approved by the ethics committee of the Faculty of Medicine at the University of Tübingen (585|2020BO) and conducted in accordance with the Declaration of Helsinki (as revised in 2013).

### Experimental procedure

The study consisted of seven laboratory visits (Fig. 1). Before the intake session (S1), participants were instructed to fast for 12 hours, with only water, unsweetened tea, or black coffee permitted. In this session, participants first answered questions regarding their mental and metabolic states (PANAS; Positive and Negative Affect Schedule, Watson et al., 1988; questions are reported in another study Fahed et al., 2024) and we drew blood (not analyzed here). In this session, participants completed a go/no-go learning task involving food and monetary rewards (∼30 min; Guitart-Masip et al., 2012; Kühnel et al., 2020), the effort allocation task (∼ 35 min, EAT; Neuser et al., 2020), and a reward rating and choice task (∼15 min). Afterwards, participants received breakfast, snacks, and monetary rewards according to task performance, and were debriefed. Participants received a smartphone to complete the Influenca app, an online gamified reinforcement learning task for repeated assessment over the following weeks (Neuser et al., 2023). Moreover, a continuous glucose sensor (Abbott Libre 3) was placed at this session (Kaduk et al., 2026). Following the first session, participants were provided with an online link to complete a questionnaire battery at home.

**Figure 1.**
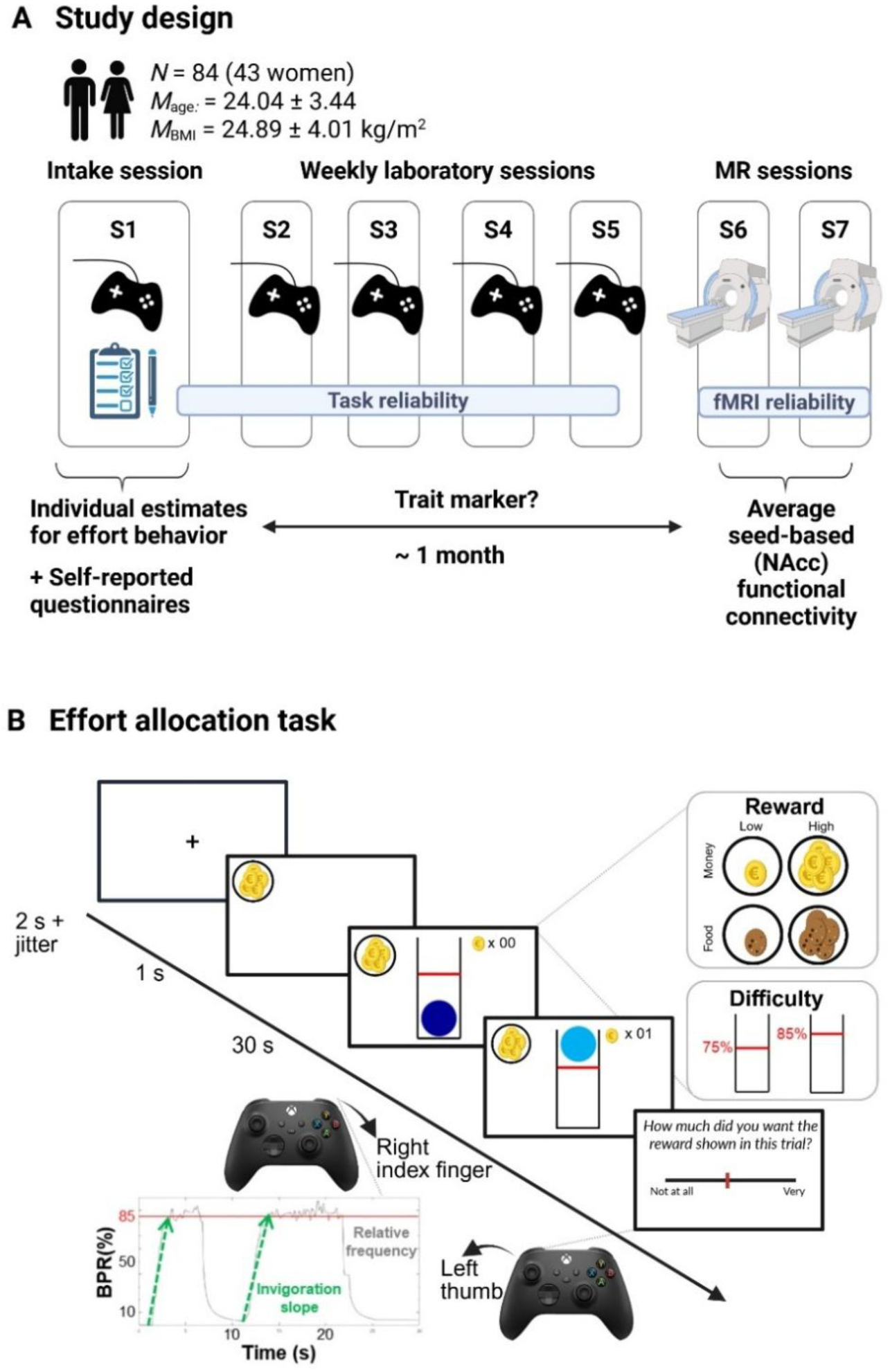
Schematic of the study and task design. A: Study design. Participants completed an intake session (S1) and four weekly behavioral sessions (S2–S5), followed by two fMRI sessions (S6–S7). Behavioral sessions assessed individual differences in effort allocation, quantified as random intercepts and slopes derived from linear mixed-effects models predicting effort maintenance (average button presses); fMRI sessions measured task-free seed-based functional connectivity (FC) from the nucleus accumbens (NAcc) to all voxels. B. Schematic of the effort allocation task. In the task, a fixation cross is shown first, followed by the reward cue. To receive a reward, participants must keep a ball above the red difficulty line (easy vs. hard) by repeatedly pressing a button with their right index finger. The magnitude of reward (low vs. high) and the type of reward (food vs. money) were additionally manipulated as task conditions. The bottom left graph shows a representative time series of a high-difficulty trial, plotting effort as button press rate (BPR) in % relative to the participant’s maximum frequency. Maintenance of effort refers to the average relative frequency (effort) during the trial. After each trial, participants used their left thumb to move a rating cursor along visual analog scales from 0 to 100, indicating how much they wanted to earn points for the presented reward and how much they had exerted themselves. The figure was adapted from (Neuser et al., 2020). Created with BioRender.com

Between the baseline session and the MRI sessions, participants attended four weekly visits (S2-S5), each lasting around 60 to 90 minutes. Within participants, sessions were scheduled at approximately the same time of day to minimize circadian variation in metabolic and hormonal state. During these sessions, participants answered questions regarding their mental and metabolic state, provided blood samples, and performed the EAT with an additional uncertainty condition (Van Den Hoek Ostende et al., 2026). To ensure task familiarity, the EAT training was repeated at the beginning of the first two weekly sessions.

The final visits of the study consisted of two MRI sessions, each lasting approximately 2.5 hours. These were typically scheduled within the same week (interval between 2 MRI sessions *M* = 2.88 d, *SD* = 2.11) and approximately 3 to 4 weeks after the last behavioral session (delay from the last behavioral session: *M* = 24.82 days, *SD* = 27.66). The timing was variable as for women, both sessions were scheduled during the luteal phase to ensure hormonal comparability across sessions (Grahlow et al., 2026). At fMRI sessions, participants arrived after an overnight fast (>12 hours) and again completed the affect and metabolic-state assessments (PANAS, Watson et al., 1988; questions are reported in another study Fahed et al., 2024) and provided blood samples before MRI acquisition. In the scanner, we collected a localizer, a structural T1-weighted image (∼6 min), a field map, a task-free scan while participants watched the Inscapes movie (Vanderwal et al., 2015) (∼10 min), and two task-based scans (∼35 min each) involving a reward learning task, once before and once after ingestion of either a milkshake or water. Here, we focus only on the task-free fMRI data that was collected in a fasting state before the manipulation of metabolic state.

### Effort allocation task

In the EAT, participants collected food and money tokens by exerting physical effort (i.e., repeatedly pressing a button with their right index finger). The task was adapted from Meyniel et al. (2013) and used frequency of button presses to measure physical effort (Neuser et al., 2020). In the intake session (S1), trials varied in reward type (food vs. money), reward magnitude (low vs. high), and difficulty (easy: 75% vs. hard: 85% of individual maximum frequency). After every effort phase, participants rated their wanting and exertion of the reward at stake on visual analog scales. Weekly sessions (S2–S5) used an adapted version with continuous difficulty levels that were either certain or uncertain (Van Den Hoek Ostende et al., 2026). Although differences in design may introduce behavioral differences compared to the first session (Table S1), coefficients were z-scored within each session to focus on the stability of relative individual differences across sessions. We therefore evaluated reliability across all sessions and additionally examined reliability across the more closely matched S2–S5 sessions.

### Questionnaires

#### Lille Apathy Rating Scale (LARS-e)

Apathy symptoms were assessed using the Lille Apathy Rating Scale (extended version; LARS-e), adapted for use in healthy populations (Bonnelle et al., 2015). The LARS-e is a 51-item self-report questionnaire assessing motivational, emotional, and behavioral engagement across four domains: action initiation, intellectual curiosity, emotional responsiveness, and self-awareness. Each item is rated on a 5-point Likert scale ranging from 1 (“completely untrue”) to 5 (“completely true”), with higher scores indicating greater motivation and engagement. Domain scores were calculated as the mean of the corresponding items, and an aggregate score was obtained by averaging the four domain means, resulting in a total score ranging from 1 to 5.

#### Snaith–Hamilton Pleasure Scale (SHAPS)

Anhedonia was measured using the Snaith–Hamilton Pleasure Scale (SHAPS), a 14-item questionnaire measuring the capacity to experience pleasure across social, sensory, food-related, and recreational domains (Snaith et al., 1995; German version: Franz et al., 1998). Items are rated on a four-point Likert scale ranging from 1 (“definitely agree”) to 4 (“strongly disagree”). Items were recoded such that higher values reflected greater anhedonia and continuous sum score was calculated.

#### Behavioral Inhibition and Activation Scales (BIS/BAS)

Behavioral inhibition and activation were recorded using the BIS/BAS scales (Carver & White, 1994; German version: Strobel et al., 2001), a 24-item questionnaire measuring sensitivity to punishment (BIS) and sensitivity to reward (BAS). Items were rated on a four-point Likert scale (1 = “very true for me” to 4 = “very false for me”), so that higher scores consistently reflected greater behavioral inhibition (BIS) or activation (BAS).

### fMRI data acquisition and preprocessing

#### Acquisition

All structural and functional MR data were acquired on a 3 T PRISMA MRI scanner using a 64-channel head coil. We first collected a structural T1-weighted image with an MP-RAGE sequence that covered the whole brain with 176 sagittal slices (flip angle = 9°, matrix size = 256 × 256, and voxel size = 1 × 1 × 1 mm³). Next, to correct for field inhomogeneities, we measured a field map using a Siemens gradient echo fieldmap sequence (short echo time (TE) = 5.19 ms and long TE = 7.65 ms, TE difference = 2.46 ms). The fMRI data were acquired using a multiband sequence (multiband factor 4) of T2*-weighted gradient echo echo-planar images. The 10-min sequence consisted of 430 volumes covering 68 axial slices and acquired with an interleaved slice order covering the whole brain, including the brain stem (repetition time (TR) = 1.4 s, TE = 30 ms, flip angle = 65°, 110 × 110 matrix, field of view = 220 × 220 mm², and voxel size = 2 × 2 × 2 mm³ (Teckentrup et al., 2021).

#### Preprocessing

Task-free fMRI data were preprocessed using fMRIPrep 20.2.7 (Esteban et al., 2019; RRID:SCR_016216), based on Nipype 1.7.0 (Gorgolewski et al., 2011; RRID:SCR_002502). Anatomical (T1-weighted) images were corrected for intensity non-uniformity, skull-stripped, segmented into gray matter, white matter, and cerebrospinal fluid, and normalized to MNI152 space using nonlinear registration. Functional images were corrected for head motion, slice-time differences, and susceptibility distortions, co-registered to the anatomical image, and normalized to MNI space (for the full fMRIPrep preprocessing workflow, see SI). Following fMRIPrep, functional data were smoothed using a 6 mm full-width-at-half-maximum Gaussian kernel in CONN (Whitfield-Gabrieli & Nieto-Castanon, 2012; RRID:SCR_009550).

#### Denoising

Functional data were denoised in CONN (v22.a) using the standard denoising pipeline (Nieto-Castanon, 2020a), including the regression of potential confounding effects of white matter signal (5 CompCor noise components), cerebro-spinal fluid signal (intermediate (default); Behzadi et al., 2007; Chai et al., 2012), motion parameters and their first order derivatives (12 factors) (Friston et al., 1996), scrubbing regressors for motion-related outlier volumes detected by Artifact Detection Tools (ART; (Nieto-Castanon, 2020b; Power et al., 2014), session, and linear trends within each functional run, followed by bandpass filtering of the BOLD time series (Hallquist et al., 2013) between 0.01 Hz and 0.1 Hz. CompCor (Behzadi et al., 2007; Chai et al., 2012) noise components within white matter and CSF were estimated by computing the average BOLD signal as well as the largest principal components orthogonal to the BOLD average, motion parameters, and outlier scans within each subject’s eroded segmentation masks.

#### First-level analysis

To estimate NAcc FC, we calculated seed-based maps in CONN using left and right NAcc seeds defined with an extended version of the Harvard-Oxford brain atlas (Desikan et al., 2006), incorporating the high-resolution, probabilistic Reinforcement Learning Atlas for subcortical nuclei (Pauli et al., 2018) and the AAL-based cerebellar regions (Tzourio-Mazoyer et al., 2002), and a nucleus tractus solitarius (NTS) mask optimized for functional imaging (Teckentrup et al., 2021; adapted from Priovoulos et al., 2019). The left and right NAcc seed-based correlation maps were then averaged, and correlation coefficients were Fisher *z*-transformed prior to second-level analyses.

### Data analysis

#### Effort allocation task

To characterize individual differences in effort-related behavior, we modeled participants’ performance in the effort allocation task from S1 using linear mixed-effects models with *lme4* for model fitting and *lmerTest* for significance testing in R, and extracted person-specific random effect coefficients.

The model to predict trial-wise relative effort includes a random intercept and slopes for reward magnitude, difficulty, reward type, their interactions, and trial number by participant (ID).

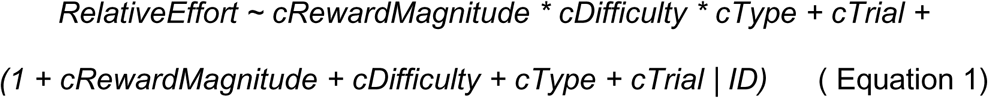

Here, cRewardMagnitude (low vs. high), cDifficulty (low vs. high), and cType (food vs. money) represent grand-mean-centered predictors; cTrial is a trial number mean-centered covariate to capture linear trends over the task. Participant-specific random effect coefficients were extracted for reward magnitude, difficulty and reward type, representing the general tendency to exert effort, reward sensitivity and cost sensitivity, respectively. These coefficients were and *z*-scored for associations with NAcc FC. To investigate associations with wanting, we repeated the procedure using wanting VAS ratings as outcome.

To evaluate the test-retest reliability of individual estimates of task behavior, we used the same linear mixed-effects model with the trial-level data of each session. Since Sessions 2-5 contained another manipulation (certain vs. uncertain trials), we only included certain trials and excluded uncertain trials to minimize differences from S1. For each session, random intercepts and slopes were extracted and *z*-scored across participants to facilitate comparison of the rank order (in addition to potential performance gains over sessions). To assess the reliability of the individual intercepts and slopes (reward magnitude and difficulty), we first calculated pairwise Pearson correlations between sessions. Second, we calculated the ICC, type consistency ICC(3,1), to quantify the stability of individual parameters across all sessions.

#### Functional connectivity and effort

To assess the test-retest reliability of NAcc FC maps across the two task-free sessions, we first examined the spatial reproducibility of group-level FC maps. Spatial reproducibility was assessed using a second-level SPM (Wellcome Department of Imaging Neuroscience, UCL) analysis with Session as a factor (along with log-transformed framewise displacement (FD), age, sex, and BMI as covariates). For each session, the resulting one-sample *t*-contrast maps were thresholded at *p* < .001 (uncorrected) and binarized. Spatial overlap between sessions was used as an index of global reliability and was quantified using the Dice and Jaccard coefficients (Dice, 1945; Jaccard, 1912; Zou et al., 2004), calculated separately for positive and negative FC.

To determine the individual reliability at the voxel- and ROI-level, we calculated the ICC(3,1) using the fmreli toolbox (Fröhner et al., 2019; beta version 1.0; https://github.com/nkroemer/reliability), with the extended Harvard-Oxford brain atlas. Voxelwise ICC maps were computed across the whole brain, and ROI-level ICC values were obtained by averaging voxelwise estimates within each atlas region. ICC values were interpreted using established cutoffs: ICC < 0.4 = poor, 0.4–0.6 = fair, 0.6–0.75 = good, > 0.75 = excellent (Shrout & Fleiss, 1979). Unthresholded statistical maps are available on NeuroVault at https://neurovault.org/collections/23461/.

#### Fingerprinting analysis

To assess whether individual NAcc-based FC patterns could re-identify participants across sessions, we used functional connectome fingerprinting based on Finn et al. (2015). One participant was excluded from the reliability analysis because they only completed one of the two scan sessions. For each participant, gray-matter-masked whole-brain NAcc seed-to-voxel FC maps were generated for the two MRI sessions, vectorized, and used as connectivity fingerprints. Re-identification was then assessed by correlating each Session 6 fingerprint with all Session 7 fingerprints using Pearson correlation, and the Session 7 map with the highest correlation was taken as the best match. Fingerprinting accuracy was defined as the proportion of participants whose best match corresponded to the same individual. To characterize the similarity structure, within-subject and between-subject correlations were Fisher *z*-transformed, and significance was evaluated using permutation testing and a binomial test against chance (Fröhner et al., 2019).

#### Second-level associations between NAcc FC and motivation measures

To associate behavioral indices of motivation with NAcc FC, we averaged FC maps from both neuroimaging sessions and used second-level regression models in SPM. We specified four separate models differing in their behavioral predictors to test how associations with NAcc FC differ across distinct motivational constructs. The first two models used person-specific random effect coefficients extracted from the EAT: random intercepts and random slopes for reward magnitude and difficulty derived from a model predicting relative effort (model 1) and subjective wanting ratings (model 2), respectively. The remaining two models used self-reported measures of motivation as predictors: scores from the LARS-e and from the SHAPS (model 3), and the BIS/BAS scales (model 4). Age, sex, BMI, and mean FD (derived from fMRIPrep) were included as covariates in all models. We explored whether the brain–behavior associations differed by sex (Fig. S1). Statistical inference was based on a voxel level threshold of *p* < .001 (uncorrected), and cluster-level significance was assessed using family-wise error (FWE) correction at *p* < .05. To facilitate visualization of significant whole-brain effects, FC estimates were extracted from the corresponding extended Harvard-Oxford brain atlas regions overlapping each significant cluster.

#### Reproducibility of brain–behavior associations

To assess the reproducibility of NAcc FC–behavior associations across behavioral sessions within the same sample, NAcc FC measures (averaged across MRI sessions S6 and S7) were related sequentially to behavioral estimates from S1 to S5. Brain– behavior associations were estimated separately using the intercept, reward-magnitude, and difficulty coefficients from each behavioral session. Across all ROIs of the Harvard-Oxford atlas, the association between NAcc FC and the behavioral coefficient was quantified using partial Pearson correlations, adjusting for age, sex, BMI, mean FD, and the other behavioral coefficients included in the corresponding model. In addition, session-wise association trajectories with all three effort parameters (intercept, reward magnitude, difficulty) were examined for the right IFG and subcallosal gyrus, corresponding to the regions identified in the original whole-brain analysis. As a sensitivity analysis, we repeated the session-specific partial correlations for the right IFG–intercept and subcallosal gyrus–difficulty slope associations, additionally adjusting for behavioral visit start time and the mean interval in days from that visit to the two MRI visits (Table S2).

To assess reproducibility across MRI sessions, the same S1 behavioral coefficients were related separately to NAcc FC from S6 and S7 using identical whole-brain models. Within each Harvard–Oxford ROI, reproducibility was quantified as the Pearson correlation between voxel-wise T-values from the two unthresholded statistical maps. This yielded one cross-session spatial correlation per ROI and behavioral effect, with higher positive values indicating more similar association patterns across MRI sessions.

#### Statistical software

Mixed-effects analyses were conducted using the *lmerTest* package (Kuznetsova et al., 2017) in R (R Core Team, 2023, version 4.4.1). EAT data were processed using MATLAB (The MathWorks, 2021b). FC analyses were performed using the CONN toolbox, release 22a (Nieto-Castanon & Whitfield-Gabrieli, 2022; Whitfield-Gabrieli & Nieto-Castanon, 2012); RRID: SCR_009550) and SPM12 (Wellcome Trust Centre for Neuroimaging, 2014; version 7771) implemented in MATLAB. Figures were generated using R, ggplot2 (Wickham, 2016) and associated packages including patchwork (Pedersen, 2024) and viridis (Garnier et al., 2021), and figures for fingerprinting were created in MATLAB (The MathWorks, 2021b). Brain– behavior association results were visualized with Mango image processing software (Lancaster et al., n.d. ; www.ric.uthscsa.edu/mango).

## Results

### Effort behavior shows high test-retest reliability, whereas NAcc FC shows spatial variability

To characterize the psychometric quality of our measures, we estimated the test-retest reliability of effort behavior and NAcc FC across sessions. As expected, reliability of motivated behavior was excellent across sessions for the intercept term (ICC = .83), and good for individual sensitivity to reward magnitude (ICC = .62) and difficulty (ICC = .68). Pairwise correlations were consistently high for the intercept (*r* = .68–.93), but more variable for reward magnitude (*r* = .29–.87) and difficulty (*r* = .44– .86). This shows that individual differences in effort behavior, particularly the general tendency to exert effort captured by the intercept, are reliable measures of motivation (Fig. 2A). For NAcc FC maps at the group-level, spatial overlap between both MRI sessions yielded overall high similarity (Jaccard: FC+ = 0.71, FC-= 0.67; Dice: FC+ = 0.83, FC-= 0.80), suggesting that group-level maps are reliable across sessions (Fig. 2B).

**Figure 2.**
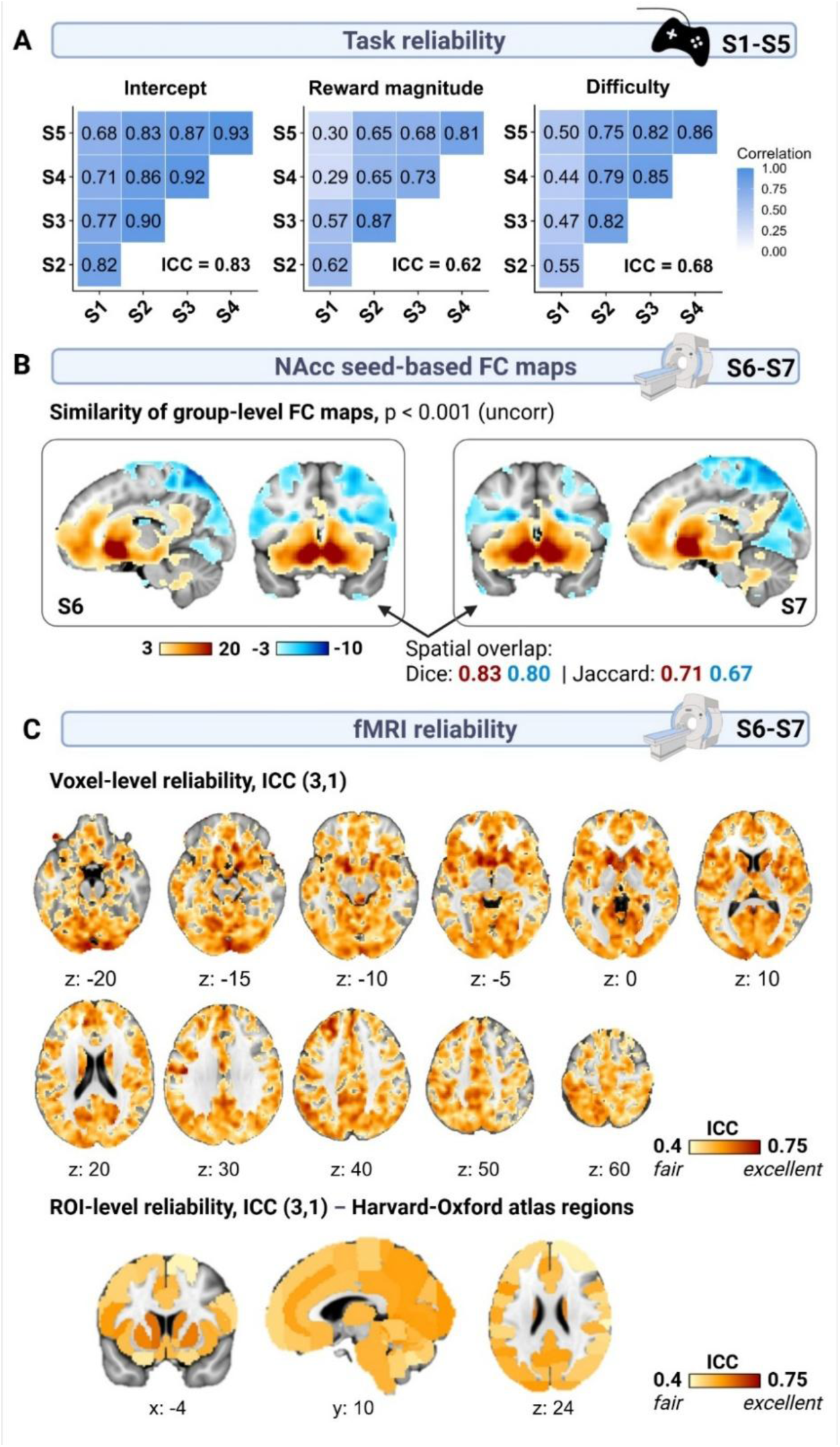
Test-retest reliability of behavioral measures of effort and NAcc functional connectivity (FC). A: Test-retest reliability of linear mixed-effects random effects coefficients across behavioral sessions revealed excellent reliability for the intercept term (ICC = 0.83), indicating stable overall motivation across time, and moderate reliability for the reward magnitude (ICC = 0.62) and reward difficulty (ICC = 0.68) effects. B: Group-level NAcc seed-based FC maps showed highly similar spatial patterns across both task-free fMRI sessions (*p* < .001, uncorrected), with strong spatial overlap for both positive (Dice = 0.83, Jaccard = 0.71) and negative (Dice = 0.80, Jaccard = 0.67) FC. C: Voxelwise test-retest reliability of NAcc seed-to-voxel FC maps show higher reliability within the core motivational circuit. Most voxels (76.72%) showed fair-to-good reliability (ICC(3,1) color scale: 0.4–0.75). ROI-level reliability estimates based on the extended Harvard-Oxford brain atlas showed mostly sufficient reliability across cortical and subcortical regions, with most regions (81.70%) showing fair-to-good reliability and only subcortical regions showing good reliability. Created with BioRender.com

At the voxel level, individual reliability was variable throughout the brain, with most voxels (76.72%) falling in the fair-to-good range (ICC .40–.75; Fig. 2C). When averaging within ROIs, 21 (13.73%) regions showed poor reliability (ICC < .40), 125 (81.70%) showed fair reliability (ICC .40–.60), and 7 (4.58%) showed good reliability (ICC > .60). Reliability was highest in subcortical regions, particularly the left ventral pallidum (ICC = .69) and right (ICC = .67) and left mammilary body (ICC = .66). These results indicate that individual-level NAcc FC is sufficiently reliable and that an average of two sessions would provide a good estimate within the core motivational network. However, the test-retest reliability was lower overall compared to task-based effort indices, specifically the intercept.

To evaluate whether FC patterns can serve as reliable individual “fingerprints” (Finn et al., 2015), we conducted identification analyses across two sessions. The reidentification rate (i.e., the percentage of participants with the highest correlation of FC maps across both fMRI sessions) was 33.73%, significantly above chance level (1/83 ≈ 1.2%; permutation test *p* < .001, 1,000 iterations). This was supported by comparing the empirical cumulative distribution functions of within-subject (*M_r_* = 0.38) and between-subjects (*M_r_* = 0.23) Fisher *z*-transformed correlations (Fig. 3). Hence, NAcc FC contains stable person-specific information that can be related to individual differences in effort-related motivation.

**Figure 3.**
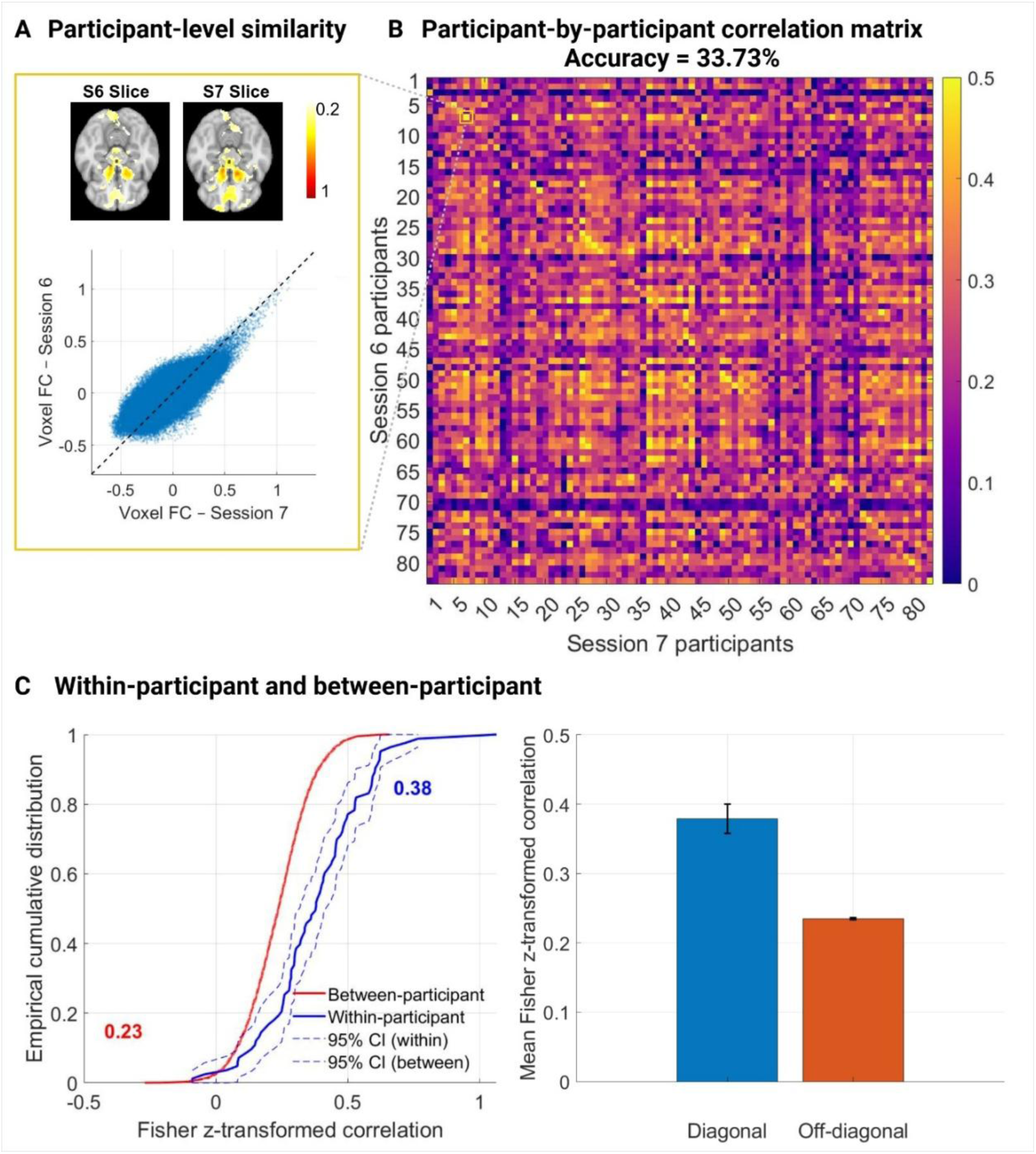
Re-identification of individuals based on nucleus accumbens (NAcc) functional connectivity (FC) profiles. A: For the participant with the highest within-participant similarity (*r* = .79), the correlation of voxelwise bilateral NAcc seed-to-voxel FC values from session 6 (S6) with the corresponding values from session 7 (S7) is shown. The upper inset shows the participant’s connectivity maps in one representative slice for both sessions, and the scatterplot below shows voxelwise FC values from S6 plotted against those from S7. The displayed value reflects the Fisher *z*-transformed Pearson correlation between this participant’s FC maps across sessions. B: Each cell represents the correlation between one participant’s connectivity map in S6 and S7. Diagonal cells reflect within-subject similarity and off-diagonal cells between-subject similarity. The yellow-outlined cell corresponds to the example participant in panel A. The accuracy indicates that 33.73% of participants were correctly identified across sessions, meaning that their connectivity pattern in S6 was most similar to their own pattern in S7. C: The left panel shows empirical cumulative distributions of Fisher *z*-transformed diagonal (within-subject, blue) and off-diagonal (between-subjects, red) correlations, with dashed lines indicating 95% confidence intervals. The right panel shows the corresponding average Fisher *z*-transformed correlations, indicating greater within-subject (*M* = 0.38) than between-subject (*M* = 0.23) similarity.

### General tendency to exert effort and sensitivity to difficulty are associated with NAcc FC

To examine whether individual differences in effort exertion are associated with NAcc FC, we included random effects of the intercept, reward magnitude, and difficulty from Session 1 linear mixed-effects model of effort maintenance as predictors for NAcc FC. The average exerted effort in the task (i.e., the intercept) was associated with increased FC between the NAcc and the right IFG (x: 46, y: 12, z: 24; *k* = 141, *p*_FWE_ = .041, *t*_max_(76) = 4.65). Since the average NAcc FC to the IFG was close to zero (*M*_NAcc-IFG FC_ = -0.073), this indicates that individuals with a greater general tendency to exert effort showed positive NAcc–IFG FC (Fig. 4A). Furthermore, more negative difficulty coefficients, reflecting stronger reductions in effort with increasing difficulty, were associated with increased FC to the subcallosal gyrus (x: -4, y: 10, z: -24; *k* = 155, *p*_FWE_ = .028, *t*_max_(76) = -5.15; Fig. 4B). Across participants, NAcc–subcallosal gyrus FC was positive on average (*M*_NAcc-subcallosal gyrus FC_ = 0.242). In other words, individuals with greater sensitivity to cost and larger reductions in effort showed higher NAcc– subcallosal connectivity. The regions identified in these analyses showed fair test-retest reliability (ICC_IFG_ = .54; ICC_subcallosal gyrus_ = .51). No clusters survived whole-brain correction for reward magnitude slopes (for unthresholded maps, see NeuroVault at https://neurovault.org/collections/23461/). The observed associations showed similar directions in women and men, with no significant sex differences (Fig. S1).

**Figure 4.**
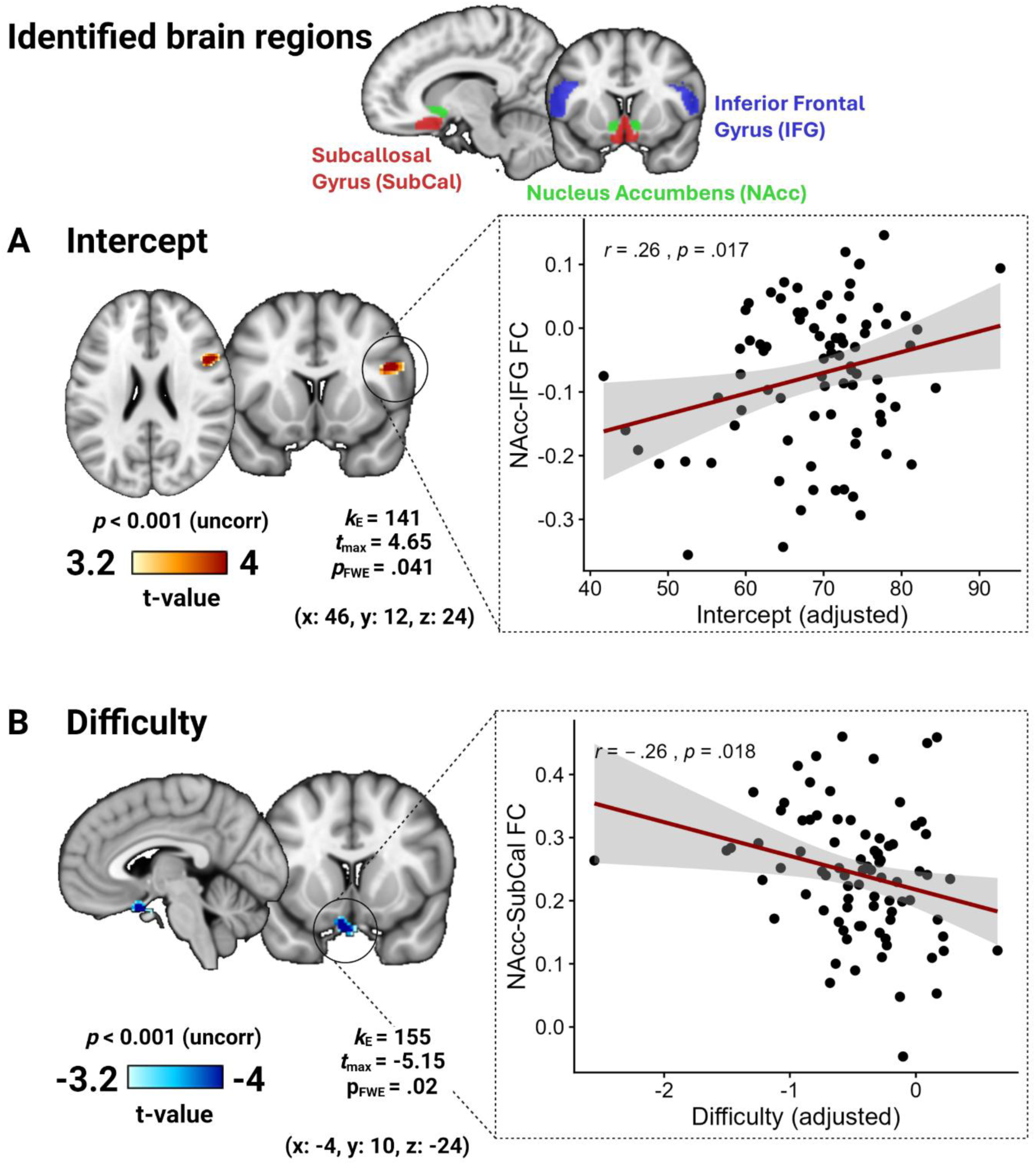
Individual estimates of task-based effort exertion are associated with nucleus accumbens (NAcc) functional connectivity (FC). A: T-value maps for the intercept (i.e., general tendency to exert effort) were voxelwise thresholded at *p* < .001 (uncorrected). Scatterplots illustrate the association between covariate-adjusted intercepts and extracted FC values (*r* = .26, *p* = .017), each controlled for age, sex, BMI, mean framewise displacement, and reward magnitude and difficulty slopes. B: T-value maps for the difficulty slopes (i.e., sensitivity to task difficulty levels) were voxelwise thresholded at *p* < .001 (uncorrected). Scatterplots illustrate the association between covariate-adjusted difficulty slope estimates and extracted FC values (*r* = -.26, *p* = .018), with both variables controlled for age, sex, BMI, mean framewise displacement, and the intercept and reward magnitude slopes. Created with BioRender.com.

### Brain–behavior association profiles vary across repeated behavioral and neuroimaging sessions

Despite the high test-retest reliability of the behavioral estimates, their ROI-wise association profiles with NAcc FC showed only moderate and heterogeneous similarity across behavioral sessions (Fig. 5A). For the intercept, similarity with S1 decreased across later sessions (S1–S2: *r* = .55; S1–S3: *r* = .33; S1–S4: *r* = .16; S1–S5: *r* = .12), and similarity among later sessions was also variable (S2–S3: *r* = .68; S3–S4: *r* = .26; S4–S5: *r* = .69). For reward magnitude, similarity was high among later sessions (S2– S3: *r* = .76; S3–S4: *r* = .81; S4–S5: *r* = .83) but low with S1 (*r*s .07–.34) and even negative between S1 and S5 (*r* = −.14). Difficulty showed an intermediate pattern, with high similarity between S1 and S2 (*r* = .63) that decreased across comparisons with later sessions (*r* = −.03 by S1–S5). Considering the two associations identified in the primary analysis (Fig. 4), the NAcc–right IFG FC association with the effort intercept was strongest in S1 (partial *r*, *r_p_* = .26) and declined toward zero by S3–S5 (S2: *r_p_* = .15; S3: *r_p_* = −.08; S4: *r_p_* = .07; S5: *r_p_* = −.01). In contrast, the NAcc–subcallosal gyrus FC association with difficulty remained negative and largely stable across all five behavioral sessions (S1:*r_p_* = −.26; S2: *r_p_* = −.18; S3: *r_p_* = −.26; S4: *r_p_* = −.24; S5: *r_p_* = −.15). Across all three behavioral effort parameters, IFG connectivity showed more variable associations across sessions, whereas the negative subcallosal gyrus– difficulty association was comparatively stable (Fig. S2).

**Figure 5.**
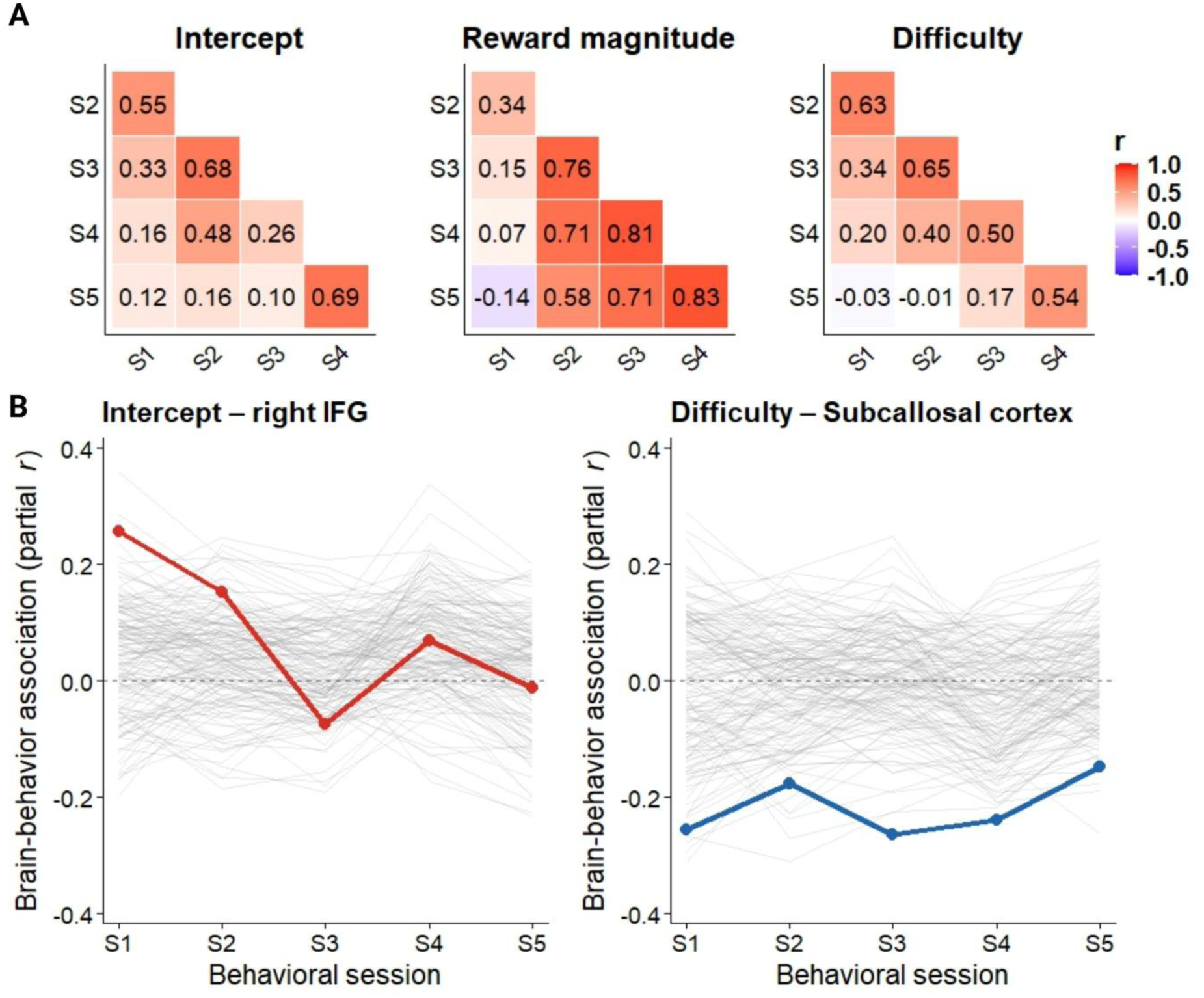
Replicability of NAcc FC–behavior associations across repeated behavioral sessions. A: Cross-session similarity of the extended Harvard–Oxford (HO) brain atlas ROI-wide brain–behavior association profiles for intercept, reward magnitude, and difficulty estimates across S1–S5. Within each ROI, associations were quantified using partial r (*r_p_*_)_, controlling for the covariates (age, sex, BMI, mean framewise displacement, behavioral estimates). Each cell shows the Pearson correlation between ROI-wise partial-r values from two behavioral sessions. B: Session-wise partial-r trajectories across all HO ROIs (gray), with the right IFG–intercept association highlighted in red and the Subcallosal–difficulty association highlighted in blue. The highlighted trajectories illustrate the stability of the two originally identified associations across repeated behavioral measurements.

Across the two MRI sessions, within ROI spatial similarity of the brain–behavior associations varied substantially among ROIs (Fig. 6A). The NAcc–right IFG intercept association showed high cross-session spatial correlation (*r* = .80), as did the NAcc– subcallosal gyrus difficulty association (*r* = .62), with both regions among the ROIs showing the highest cross-session spatial reproducibility. Across behavioral estimates and sessions, ROI-level FC reliability (ICC) was not consistently related to the strength of brain–behavior associations (correlations computed separately for each behavioral session S1–S5; *r*_intercept_ = −.21 to .14; *r*_rewmag_ = −.14 to .30; *r*_difficulty_ = −.02 to .45, Fig. S3).

**Figure 6.**
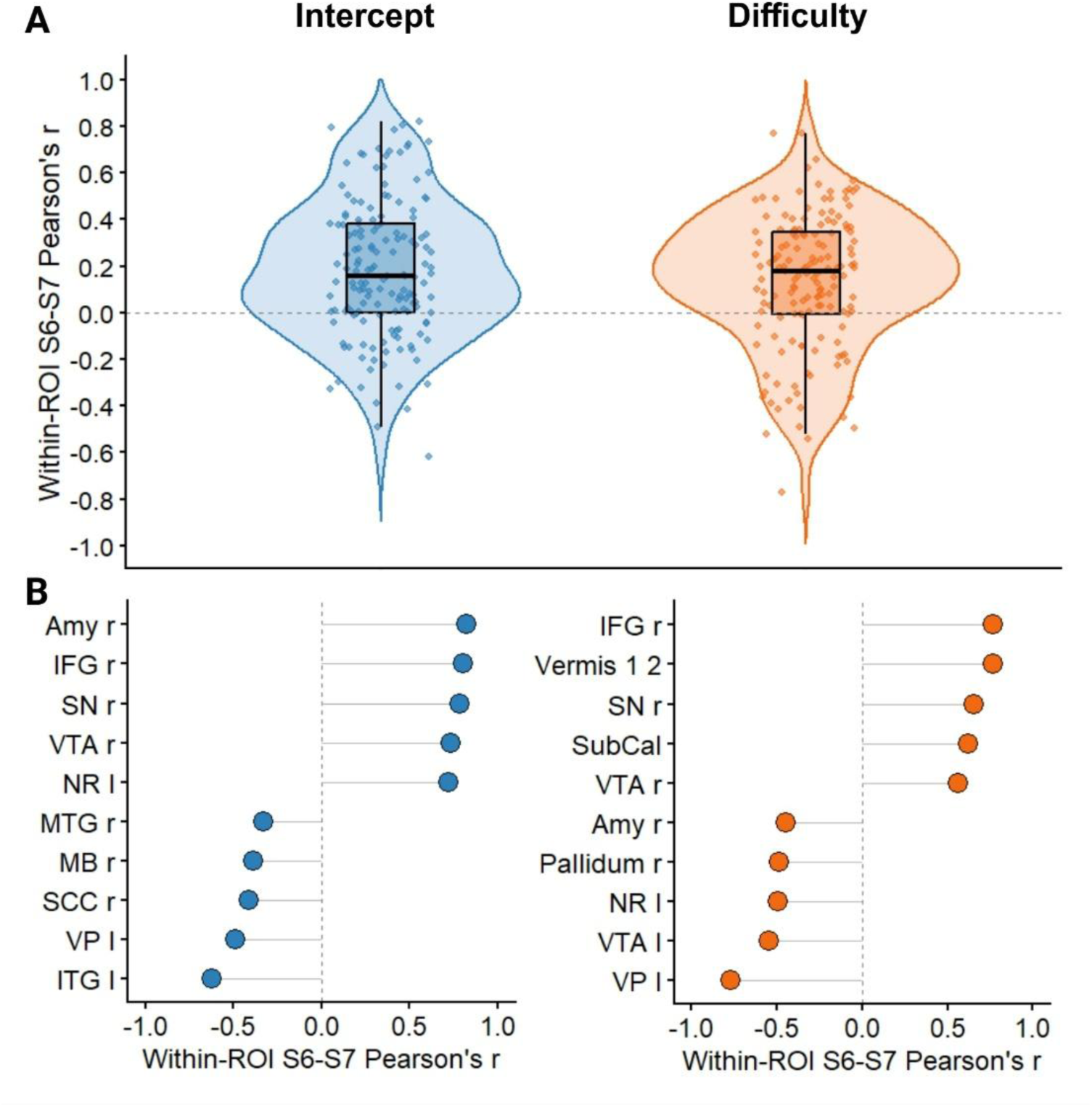
Cross-session spatial similarity of region of interest (ROI)-wise behavioral association maps. A: Violin plots showing the distribution of within-ROI spatial Pearson correlations between Session 6 (S6) and Session 7 (S7) unthresholded *T*-values for the intercept and difficulty slopes across the extended Harvard–Oxford brain atlas ROIs. Points represent individual ROIs and boxplots show the median and interquartile range. B: Five ROIs with the highest and five with the lowest spatial correlations for intercept and difficulty coefficients as predictors. Positive values indicate similar voxel-wise association patterns across sessions, whereas negative values indicate inverse spatial patterns. ROI: region of interest; r: right; l: left; Amy: amygdala; IFG: inferior frontal gyrus; SN: substantia nigra; VTA: ventral tegmental area; MTG: middle temporal gyrus; MB, midbrain; SCC: supracalcarine cortex; VP: ventral pallidum; IT: inferior temporal gyrus; SubCal: subcallosal gyrus; NR: nucleus ruber.

### Relationship between behavioral effort, subjective wanting and self-reported motivational trait measures

To examine how behavioral effort relates to other motivational measures, we assessed associations with subjective wanting ratings and self-reported motivational traits. Behavioral effort indices were strongly correlated with wanting rating indices (reward magnitude: *r* = .79; difficulty: *r* = .56; intercept: *r* = .44; all *p* < .001; Fig. S4). In contrast, behavioral effort indices showed weak associations with BIS/BAS questionnaire scores (i.e., the highest association between BAS-difficulty: *r* = .24, *p* = .027), and non-significant correlations for SHAPS and LARS-e (all *r*s < .25, *p* > .055; Fig. S5 and Fig. S6). To assess whether the observed NAcc FC associations were specific to behavioral effort indices vs. self-report measures of motivation, we tested whether wanting-related coefficients (intercept, reward magnitude, and difficulty slopes) and self-reported questionnaire measures (LARS-e apathy scores, SHAPS anhedonia scores, and BIS/BAS scores) are associated with NAcc FC. In contrast to the behavioral effort indices, none of these alternative motivational measures yielded clusters surviving whole-brain correction at *p*_FWE_cluster_ = .05 (Table S3).

## Discussion

Effort exertion is thought to be supported by the recruitment of a motivational brain circuit, which is functionally indexed with seed-based FC to the NAcc as the major hub (Salamone & Correa, 2012). Here, we investigated whether trait-like individual differences in effort-based decision-making are reflected in intrinsic FC of the NAcc at rest, and whether such brain–behavior associations are reproducible over repeated behavioral and fMRI measurements. As expected, effort-based behavior on the EAT (Van Den Hoek Ostende et al., 2021) showed high test–retest reliability. In contrast, NAcc FC showed spatially heterogeneous reliability with sufficient individual-specific information across sessions demonstrated by a decent re-identification accuracy based on FC profiles. Behavioral differences in effort allocation were associated with NAcc connectivity to the right IFG (overall exertion) and the subcallosal gyrus (difficulty slopes). Repeated behavioral measurements further revealed distinct reproducibility profiles: the IFG association decreased over behavioral assessments, whereas the subcallosal association was stable. Among ROIs, both showed comparatively high spatial reproducibility across MRI sessions, suggesting that they are suitable targets for brain–behavior associations with effort. In contrast to behavioral indices, wanting ratings and self-reported questionnaire measures of motivation did not yield comparable corrected associations. Together, these findings show that reliable behavioral measures do not guarantee equally reproducible brain–behavior associations, and that repeated assessments can reveal which associations are stable and which are session-dependent.

In line with our expectations, NAcc FC at rest was associated with reliable individual differences in effort. Individuals exerting more effort overall showed stronger positive FC between the NAcc and the right IFG. The right IFG helps initiate inhibitory control (Choo et al., 2022) and responds to cues to continue an action (Wang, 2025). Its posterior portion also contributes to adjusting planned movements (Maizey et al., 2020), which may be relevant when effortful behavior requires responses to changing task demands. Consistent with a motivational role, lower connectivity between the NAcc core and an IFG/insula region has been linked to more severe anticipatory anhedonia in depression (Hu et al., 2023). In contrast, greater cost sensitivity as indexed by negative slopes with increasing task difficulty was associated with higher NAcc–subcallosal FC.

Studies in marmosets have shown that overactivation of subcallosal area 25 reduced the willingness to respond for a reward as the required effort increased (Alexander et al., 2019), whereas inactivation of the area increased their willingness to respond (Banai Tizkar et al., 2024). Activating projections from area 25 to the NAcc also blunted anticipatory responses to reward cues (Wood et al., 2023). In humans, lesions involving area 25 were associated with greater effort discounting when participants worked for another person relative to themselves (Lockwood et al., 2024). Liu et al. (2021) also linked the subgenual region to consummatory pleasure, finding that the association between pleasure ratings and right NAcc shell– subgenual/pregenual ACC connectivity differed between patients with depression and healthy controls. Higher NAcc–subcallosal connectivity may therefore reflect stronger coupling between affective valuation and motivational processes, potentially contributing to greater adjustment of effort as difficulty increases.

The repeated behavioral assessments revealed different trajectories of brain– behavior associations. The NAcc-IFG association was associated with the initial tendency to exert effort, but this relationship weakened over time. In contrast, the NAcc-subcallosal gyrus association with difficulty showed stable associations with repeated assessments of the task. This may reflect meaningful session-dependent variation in the behavioral expression of motivation (Racicot et al., 2024). It is conceivable that participants adjust their behavior over time and become more strategic in investing their effort, which may explain the gradual decrease in reliability of the intercept coefficient. In other words, early effort exertion may be driven by other processes, such as novelty seeking (Gruber et al., 2017), compared to sustained and habitual effort exertion (Yang et al., 2022). Instead, effort costs have been previously shown to be trait-like (Church & Alsop, 2026; Mehrhof & Nord, 2025), and this could be ingrained in individual differences in the functional architecture of the motivational circuit. Together, these findings highlight the value of repeated behavioral assessments beyond measuring test-retest reliability since the trajectories of brain– behavior associations may improve their interpretation.

Spatial similarities of brain–behavior association patterns varied considerably across regions, yet both the NAcc–IFG association with general effort tendency and the NAcc–subcallosal gyrus association with difficulty sensitivity showed comparatively high similarity across MRI sessions. Particularly for the NAcc-IFG FC, this contrasted with its changing behavioral correspondence across assessments, suggesting that reliability of the brain–behavior association is primarily due to the behavioral changes over time. Regional reliability of NAcc-based FC also showed no clear relationship with brain–behavior associations, consistent with evidence that reliability is necessary, but not sufficient to predict behavior (Noble et al., 2017, 2019). These findings complement recent work showing that brain–behavior associations are often small-to-medium, sample-dependent, and sensitive to measurement reliability (Gell et al., 2024; Kang et al., 2024; Marek et al., 2022). Repeated measurements of both behavior and fMRI provide a complementary aspect by testing whether a brain–behavior association is stable when the same individuals are assessed again. Since repeating behavioral assessments is potentially more cost-effective (De Haas, 2018) compared to repeated fMRI sessions, this may provide a scalable solution to improve research on brain– behavior associations in medium-sized samples (Spisak et al., 2023).

Beyond brain–behavior associations, effort-based behavior showed high test– retest reliability across repeated assessments, extending previous evidence that effort-based tasks capture meaningful individual differences in motivated behavior (Malik et al., 2025; Ohmann et al., 2022; Van Den Hoek Ostende et al., 2021). Also, NAcc FC patterns showed high spatial overlap, even if reliability was mixed at the voxel level. Perhaps surprisingly, the reliability of NAcc seed-based FC has received little attention in the context of individual-differences research on motivation, and the present estimates may serve as a useful benchmark for future work, including for translational research (Kroemer et al., 2022; H. Li et al., 2026; Liu et al., 2021; Morris et al., 2023).

In contrast to effort indices, wanting ratings and self-reported motivational measures did not show comparable corrected NAcc FC associations. This is consistent with evidence that behavioral and self-report measures capture partly distinct aspects of motivation (Dang et al., 2020; Friedman & Banich, 2019; Renz et al., 2023), which are reflected in non-overlapping neural correlates (Bonnelle et al., 2015; Derosiere et al., 2025). Our findings therefore suggest that motivational questionnaires tap into separable behavior and neural processes, suggesting that they cannot replace behavioral assessment as proxy measures for the same targets (e.g., to enhance effort with circuit-based interventions).

### Limitations

Despite the strengths of the work, such as the many repeated behavioral sessions and two independent fMRI runs, several limitations should be considered. First, behavioral and neuroimaging measures were sampled with different resolution (behavior: 5 sessions; task-free FC: 2 sessions). The behavioral sessions S2-S5 were also not scheduled in the morning, and the interval between behavioral and neuroimaging assessments was variable. However, adjusting for behavioral visit start time and the mean interval to the two MRI visits had a negligible effect on the associations, changing all ten session-specific partial correlations by less than .03 and preserving the pattern across sessions (Table S2). Second, FC reliability varies across regions, scan length, and preprocessing choices, with subcortical connections particularly susceptible to these factors (Noble et al., 2021; Shah et al., 2016). Nevertheless, decent re-identification rates based on individual NAcc FC patterns indicate that they retain substantial individual-specific information, which is necessary for prediction of individual differences (Fröhner et al., 2019). Third, the healthy sample likely restricted variation in apathy and anhedonia, reducing sensitivity for some questionnaire-based associations. Still, characterizing motivated behavior in a non-clinical sample is valuable as a general benchmark, and associations with other questionnaires intended for healthy participants suggest that this is unlikely to explain the weak associations. Fourth, physiological state factors were only partially controlled. All task-free scans were acquired in the luteal phase in naturally cycling women and participants were fasted and went to the scanner at approximately the same time of day, which likely reduced state-related variability. However, by design, behavioral effort was sampled across the menstrual cycle, and motivation varies across cycle phases (Grahlow et al., 2026). Nevertheless, stable between-person differences in motivation can be detected despite systematic within-person fluctuations (Hewitt et al., 2025), and the high test-retest reliability shows that state variability is low compared to trait-like differences in behavior.

### Conclusion

Behavioral differences in effort were reliable across repeated assessments and associated with NAcc FC at rest. Overall effort exertion was associated with NAcc–IFG FC, but this association declined for later behavioral sessions. In contrast, effort cost sensitivity was associated with NAcc–subcallosal FC and this association was stable over repeated behavioral sessions. Our findings show that trait-like individual differences in behavior do not necessarily translate into equally reproducible neural associations and underscore the importance of evaluating the reproducibility of brain– behavior relationships in addition to the psychometric quality of the measures themselves. Crucially, by evaluating changes in brain–behavior associations over repeated sessions of an effort task, it might be possible to derive more mechanistic insights about the functional role of such FC findings.

## Acknowledgement

We thank Antonia Schlaich, Johanna Voß, Franziska Peglow, Hannah Groß, Sophie Mathis, David Bartsch, Rauda Fahed, Katharina Bertl, Johanna Theuer, Marie Kaeber, Ebru Sarmisak, Laura Heidiri, and Ricarda Bode for help with data acquisition, and Asma Ikhenazene and Alica Guzman for support in processing the functional connectivity data. The study was supported by the German Research Foundation (DFG), grants DE 2319/22-1; KR 4555/7-1; KR 4555/9-1; KR 4555/10-1. ZsK was supported by the International Research Training Group: Women’s Mental Health Across the Reproductive Years (IRTG 2804), funded by the DFG; grant number: GRK 2804/1.

## Author contributions

NBK and BD were responsible for the study concept and design. MG, KK, & AK collected data under supervision by NBK and BD. NBK conceived the method of this paper, and AK preprocessed the data. ZsK, CS & NBK performed the data analysis and prepared the figures AK contributed to the analyses. ZsK & NBK wrote the manuscript. All authors contributed to the interpretation of findings, provided critical revision of the manuscript for important intellectual content and approved the final version for publication.

