## Supplementary material for "How reliable is the link between effort and nucleus accumbens connectivity at rest? A psychometric evaluation of repeated assessments": Table S1

### Supplementary Material A: fMRI preprocessing

#### Resting-state fMRI preprocessing with fMRIPrep

The full preprocessing workflow for resting-state fMRI data was carried out with fMRIPrep 20.2.7 [@fmriprep1; @fmriprep2; RRID:SCR_016216], based on Nipype 1.7.0 [@nipype1; @nipype2; RRID:SCR_002502]. The pipeline details summarized below expand on the preprocessing steps applied prior to smoothing in the CONN toolbox as described in the main text.

##### *Anatomical preprocessing*

A total of two T1-weighted (T1w) images were found within the input BIDS dataset. All T1w images were corrected for intensity non-uniformity with N4BiasFieldCorrection (from ANTs 2.3.3; Tustison et al., 2010). The T1w reference was skull-stripped with a Nipype implementation of the antsBrainExtraction.sh workflow (ANTs), using the OASIS30ANTs template as a target. Brain tissue segmentation of cerebrospinal fluid (CSF), white matter (WM), and gray matter (GM) was performed on the brain-extracted T1w using fast from FSL 5.0.9 (Zhang et al., 2001). A T1w reference image was computed after registration of the two T1w images (following INU correction) using mri_robust_template (FreeSurfer 6.0.1; Reuter et al., 2012). Brain surfaces were reconstructed using recon-all (FreeSurfer 6.0.1; Fischl et al., 1999), and the brain mask estimated previously was refined with a custom variation of the method to reconcile ANTs-derived and FreeSurfer-derived segmentations of the cortical gray matter of Mindboggle (Klein et al., 2017). Volume-based spatial normalization to the MNI152NLin2009cAsym template was performed through nonlinear registration with antsRegistration (ANTs 2.3.3; Avants et al., 2011). The selected template for spatial normalization was the ICBM 152 Nonlinear Asymmetrical template version 2009c (Fonov et al., 2009, 2011).

##### *Functional preprocessing*

For each of the 11 BOLD runs per subject (across tasks and sessions), the following preprocessing steps were performed. A reference volume and its skull-stripped version were generated by aligning and averaging the single-band reference (SBRef) image. A B0-nonuniformity map (or fieldmap) was estimated based on a phase-difference map calculated from a dual-echo GRE (gradient-echo) sequence, processed with a custom workflow of SDCFlows (Roebuck et al., 2021) inspired by the epidewarp.fsl script and improvements in the HCP Pipelines (Glasser et al., 2013). The fieldmap was co-registered to the target EPI reference run and converted to a displacement field map using FSL's fugue and other SDCflows tools. Using the estimated susceptibility distortion, a corrected EPI reference was calculated to enable more accurate co-registration with the anatomical reference. The BOLD reference was then co-registered to the T1w reference using bbregister (FreeSurfer; Greve & Fischl, 2009), configured with six degrees of freedom. Head-motion parameters with respect to the BOLD reference (transformation matrices and six corresponding rotation and translation parameters) were estimated before any spatiotemporal filtering using mcflirt (FSL 5.0.9; Jenkinson et al., 2012). BOLD runs were slice-time corrected to the midpoint of the slice acquisition range (0.65 s, corresponding to 0 s–1.3 s) using 3dTshift from AFNI 20160207 (Cox, 1996). The BOLD time-series (including slice-time correction where applied) were resampled onto their original native space by applying a single composite transform to correct for head motion and susceptibility distortions. These resampled BOLD time-series are referred to as preprocessed BOLD in native space (or simply preprocessed BOLD). The same time-series were then resampled into standard space (MNI152NLin2009cAsym), generating preprocessed BOLD runs in MNI152NLin2009cAsym space. Several confounding time-series were calculated from the preprocessed BOLD: framewise displacement (FD), DVARS, and three region-wise global signals extracted from CSF, WM, and the whole-brain masks. FD was computed using two formulations following Power et al. (2012) and Jenkinson et al. (2012). The three global signals were extracted within the respective masks. Physiological regressors were extracted for component-based noise correction (CompCor; Behzadi et al., 2007). Principal components were estimated after high-pass filtering the preprocessed BOLD time-series with a discrete cosine filter with a 128 s cut-off. For temporal CompCor (tCompCor), components were calculated from the top 2% of most variable voxels within the brain mask. For anatomical CompCor (aCompCor), three probabilistic masks (CSF, WM, and combined CSF+WM) were generated in anatomical space. The implementation differs from Behzadi et al. (2007) in that the aCompCor masks were subtracted from a mask of voxels likely to contain a minimal fraction of GM, obtained by dilating a GM mask extracted from FreeSurfer's aseg segmentation. These masks were then resampled into BOLD space and binarized by thresholding at 0.99. Components were also calculated separately within the WM and CSF masks. For each CompCor decomposition, the k components with the largest singular values were retained such that the retained components explain 50% of variance across the nuisance mask (CSF, WM, combined, or temporal). The remaining components were dropped from consideration. The head-motion estimates were included in the corresponding confounds file. The confound time-series derived from head motion estimates and global signals were expanded with temporal derivatives and quadratic terms for each regressor (Satterthwaite et al., 2013). Frames exceeding a threshold of 0.5 mm FD or 1.5 standardized DVARS were annotated as motion outliers. All resamplings were performed in a single interpolation step by composing all relevant transformations (head-motion matrices, susceptibility distortion correction when available, and co-registrations to anatomical and output spaces). Gridded (volumetric) resamplings used antsApplyTransforms (ANTs), configured with Lanczos interpolation to minimize smoothing effects of other kernels (Lanczos, 1956). Non-gridded (surface) resamplings used mri_vol2surf (FreeSurfer).

Many internal operations of fMRIPrep utilize Nilearn 0.6.2 (Abraham et al., 2014; RRID:SCR_001362), particularly within the functional processing workflow. For further details on the pipeline, see the workflows section of the fMRIPrep documentation (https://fmriprep.readthedocs.io/en/latest/workflows.html).

### Supplementary Material B: Influence of the first session on effort reliability

**Table S1. Sensitivity analysis of test-retest reliability of effort indices, with and without Session 1 (S1).**

| Parameter | ICC (S1–S5) | ICC (S2–S5) |
|---|---|---|
| Intercept | 0.830 | 0.886 |
| Reward magnitude | 0.618 | 0.732 |
| Difficulty | 0.684 | 0.817 |

*Note*: ICC(3,1) consistency estimates computed on values *z*-scored within session. n = 93 participants with complete data across all five sessions.

### Supplementary Material C: Exploratory analysis of sex differences in brain–behavior associations

To examine whether the identified brain–behavior associations differed between women and men, we conducted exploratory sensitivity analyses including interactions between sex and the respective behavioral parameters. We tested the sex × intercept interaction for NAcc–right IFG functional connectivity and the sex × difficulty interaction for NAcc–subcallosal gyrus connectivity. Models were adjusted for age, BMI, mean framewise displacement, and the other behavioral coefficients.

Both associations showed comparable directions and slopes in women and men (Fig. S1). Neither the sex × intercept interaction ($\beta$ = 0.001, 95% CI [−0.051, 0.052], $p$ = .978) nor the sex × difficulty interaction ($\beta$ = 0.003, 95% CI [−0.046, 0.051], $p$ = .908) was significant. Thus, these exploratory analyses provided no evidence that the identified brain–behavior associations differed by sex, although the absence of significant interactions does not establish equivalence between groups.

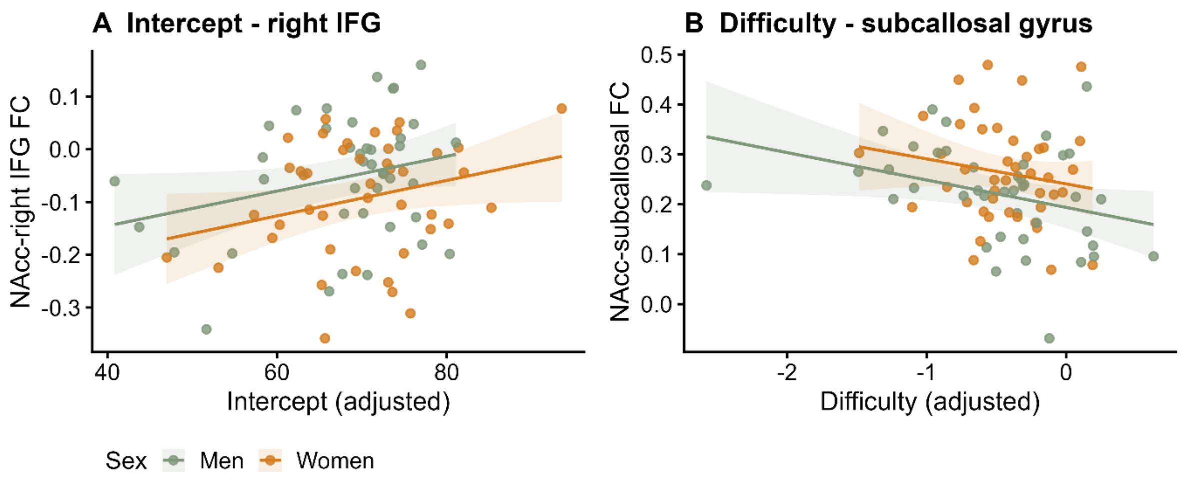


**Figure S1. Exploratory analysis of NAcc FC–behavior associations by sex.** A: Scatterplots illustrate the association between covariate-adjusted intercept estimates (i.e., general tendency to exert effort) and extracted NAcc–right IFG FC values, shown separately for women (orange) and men (green), with both variables adjusted for age, BMI, mean framewise displacement, and reward magnitude and difficulty slopes. B: Scatterplots illustrate the association between covariate-adjusted difficulty slope estimates (i.e., sensitivity to task difficulty levels) and extracted NAcc–subcallosal gyrus FC values, shown separately for women (orange) and men (green), with both variables adjusted for age, BMI, mean framewise displacement, and the intercept and reward magnitude slopes. Lines represent sex-specific fitted associations with 95% confidence intervals. Exploratory sex-by-behavior interactions were tested using multiple regression models including the corresponding covariates.

### Supplementary Material D: Assessment timing and brain–behavior associations

**Table S2. Sensitivity of NAcc connectivity–effort behavior associations to assessment timing.**

| FC–behavior association | Session | Original partial *r* | Timing-adjusted partial *r* | Δr |
|---|---|---|---|---|
| NAcc–right IFG / Intercept | S1 | 0.257 | 0.281 | +0.024 |
| | S2 | 0.152 | 0.178 | +0.026 |
| | S3 | −0.075 | −0.057 | +0.018 |
| | S4 | 0.071 | 0.099 | +0.027 |
| | S5 | −0.012 | −0.004 | +0.008 |
| NAcc–subcallosal gyrus / Difficulty | S1 | −0.254 | −0.233 | +0.021 |
| | S2 | −0.173 | −0.153 | +0.020 |
| | S3 | −0.263 | −0.238 | +0.025 |
| | S4 | −0.239 | −0.210 | +0.029 |
| | S5 | −0.148 | −0.132 | +0.016 |

*Note:* $N$ = 83 for every comparison. Both models controlled for age, sex, BMI, framewise displacement, and the other two behavioral coefficients. The timing-adjusted model additionally controlled for behavioral visit start time and the mean absolute interval (days) from the behavioral session to the two MRI visits. Δr is the adjusted minus the original partial correlation, calculated before rounding. NAcc: nucleus accumbens; IFG: inferior frontal gyrus; FC: functional connectivity.

### Supplementary Material E: Brain–behavior associations across behavioral sessions

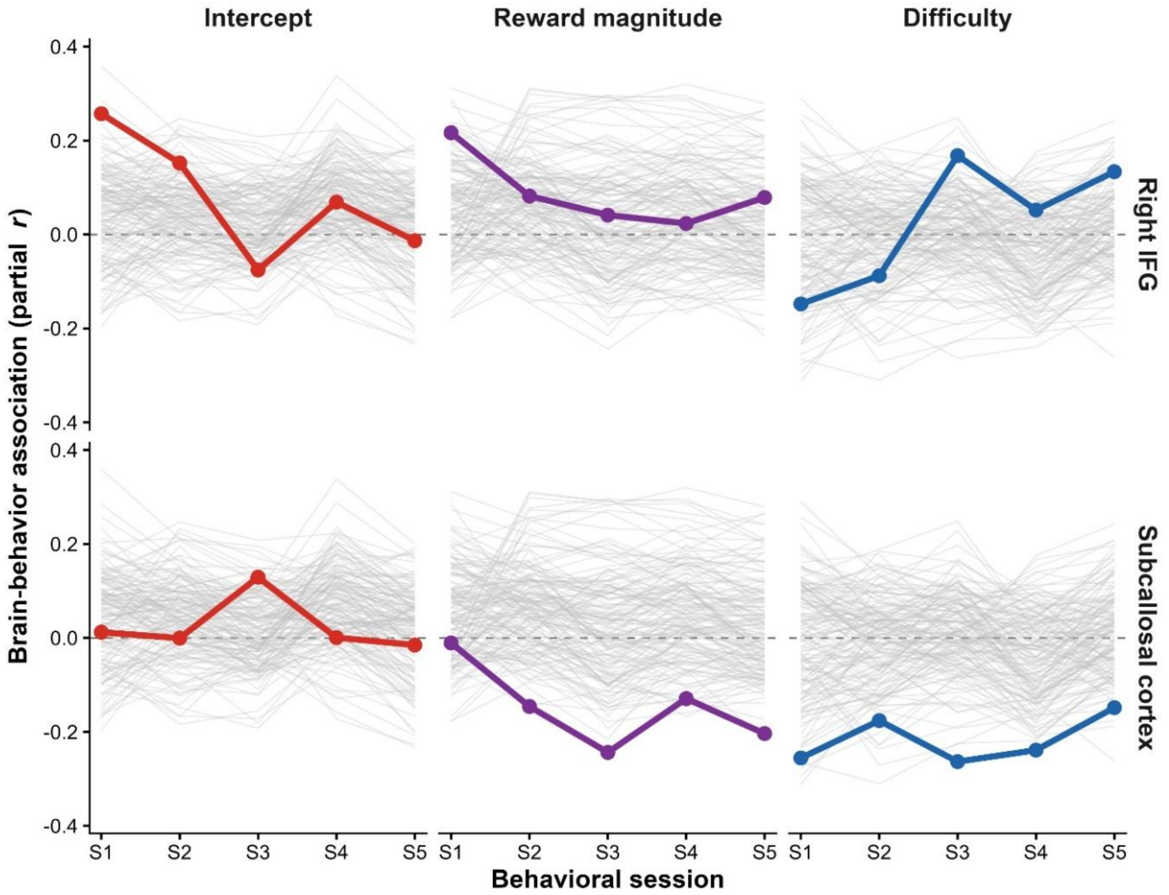


**Figure S2. Session-wise NAcc FC–behavior associations across regions and functional systems.** Session-wise brain–behavior associations across behavioral sessions S1–S5 for the intercept, reward magnitude, and difficulty estimates. Gray lines show partial correlations for individual Harvard–Oxford ROIs. Colored lines highlight associations for the right inferior frontal gyrus (IFG; upper row) and subcallosal cortex (lower row), corresponding to the regions identified in the primary whole-brain analysis. Partial correlations were adjusted for age, sex, BMI, mean framewise displacement, and the other behavioral estimates included in the corresponding model.

### Supplementary Material F: Functional connectivity reliability and brain–behavior association

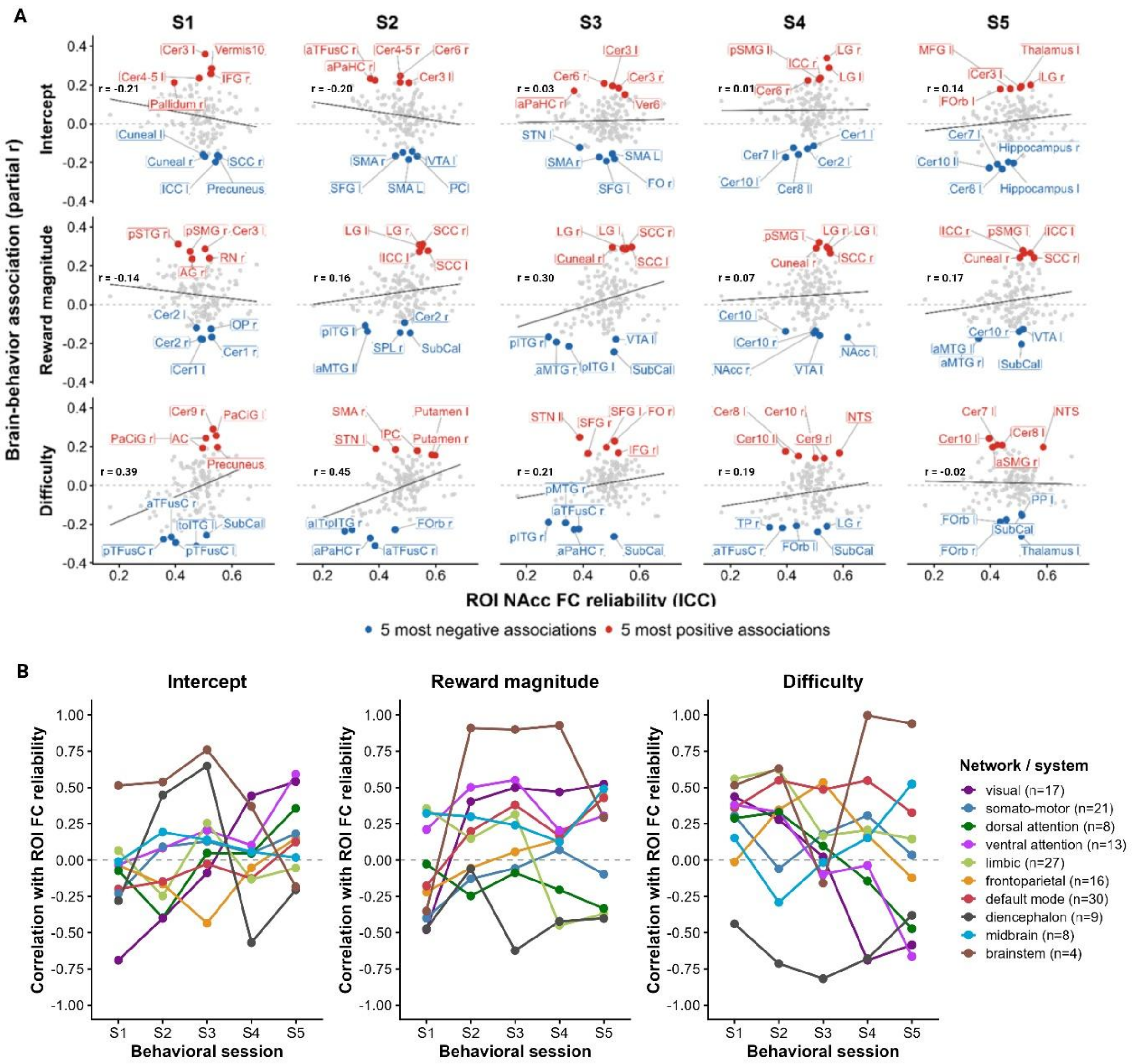

**Figure S3. Relationship between ROI-level NAcc FC reliability and brain–behavior associations across behavioral sessions.** A: Scatterplots show the relationship between ROI-level NAcc functional connectivity (FC) test–retest reliability and ROI-wise brain–behavior associations for the intercept, reward magnitude, and difficulty estimates across behavioral sessions S1–S5. Each point represents one Harvard–Oxford ROI. The x-axis shows the reliability of NAcc–ROI FC across MRI sessions S6 and S7 (ICC), and the y-axis shows the corresponding partial correlation between NAcc FC and the behavioral estimate, adjusted for age, sex, BMI, mean framewise displacement, and the other behavioral estimates included in the corresponding model. Black lines show the linear relationship across all included ROIs, and the Pearson correlation coefficient for this relationship is shown in each panel. The five most positive and five most negative brain–behavior associations within each panel are highlighted in red and blue, respectively, and labeled by ROI. B: Network-wise relationship between NAcc FC reliability and brain–behavior associations. For each behavioral estimate and session, points show the Pearson correlation, across ROIs within each functional network, between NAcc FC test–retest reliability across S6 and S7 and the corresponding ROI-wise partial brain–behavior correlations. Lines connect estimates across behavioral sessions. The number of ROIs included in each network/system is indicated in the legend.

### Supplementary Material G: Behavioral effort estimates associations with wanting estimates, and questionnaire scores

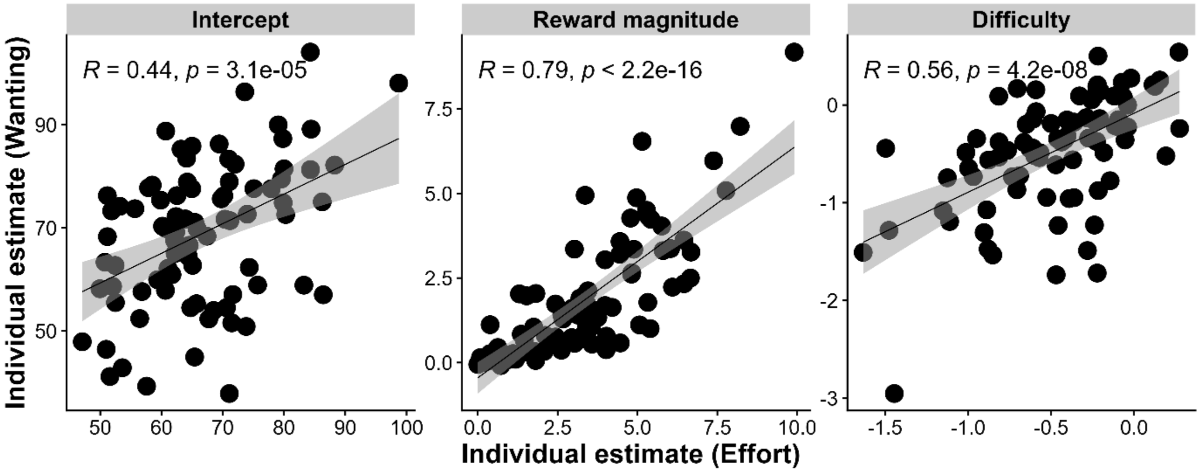


**Figure S4. Relationship between behavioral effort and wanting estimates.** Each panel shows the association between behavioral effort (x-axis) and wanting (y-axis) coefficients for intercept (left), reward magnitude (center), and difficulty (right). Points represent participants, and regression lines with 95% confidence intervals are shown. All correlations are positive and significant (*r* = .44–.79, *p* < .001), indicating that greater effort exertion was associated with higher wanting ratings across conditions.

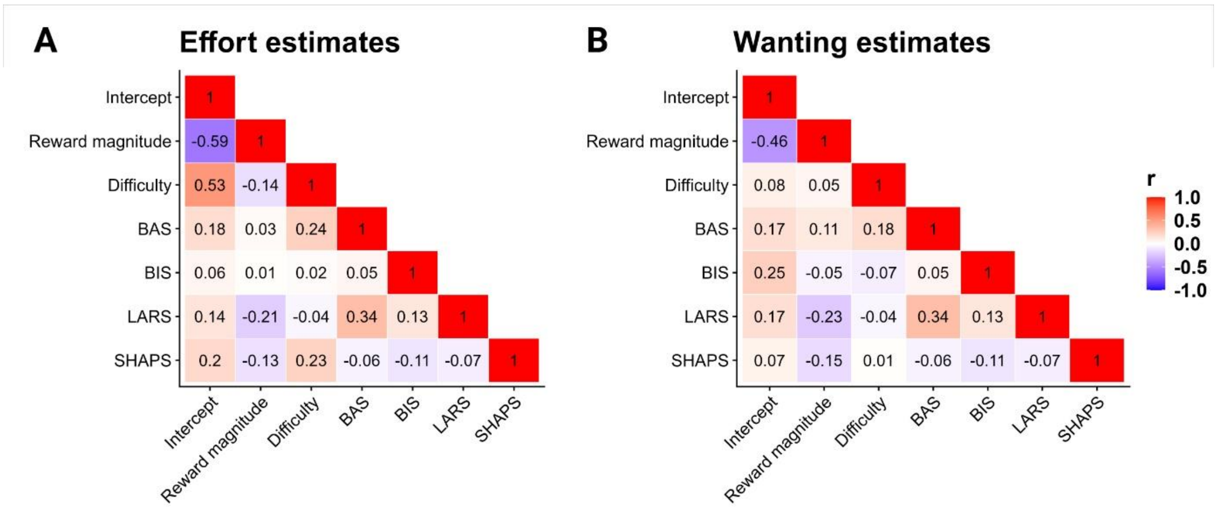


**Figure S5. Correlation matrix of behavioral effort and wanting estimates.** A: Heatmap showing pairwise correlations between behavioral effort indices (intercept, reward magnitude, difficulty) and questionnaire measures (BAS, BIS, LARS-e, SHAPS). B: Heatmap showing pairwise correlations between wanting estimates (intercept, reward magnitude, difficulty) and questionnaire measures (BAS, BIS, LARS-e, SHAPS). Red indicates positive correlations, blue negative; intensity reflects magnitude. The strongest correlations are between intercepts and reward magnitude slopes (*r* = -.59) and intercepts and difficulty slopes (*r* = .53).

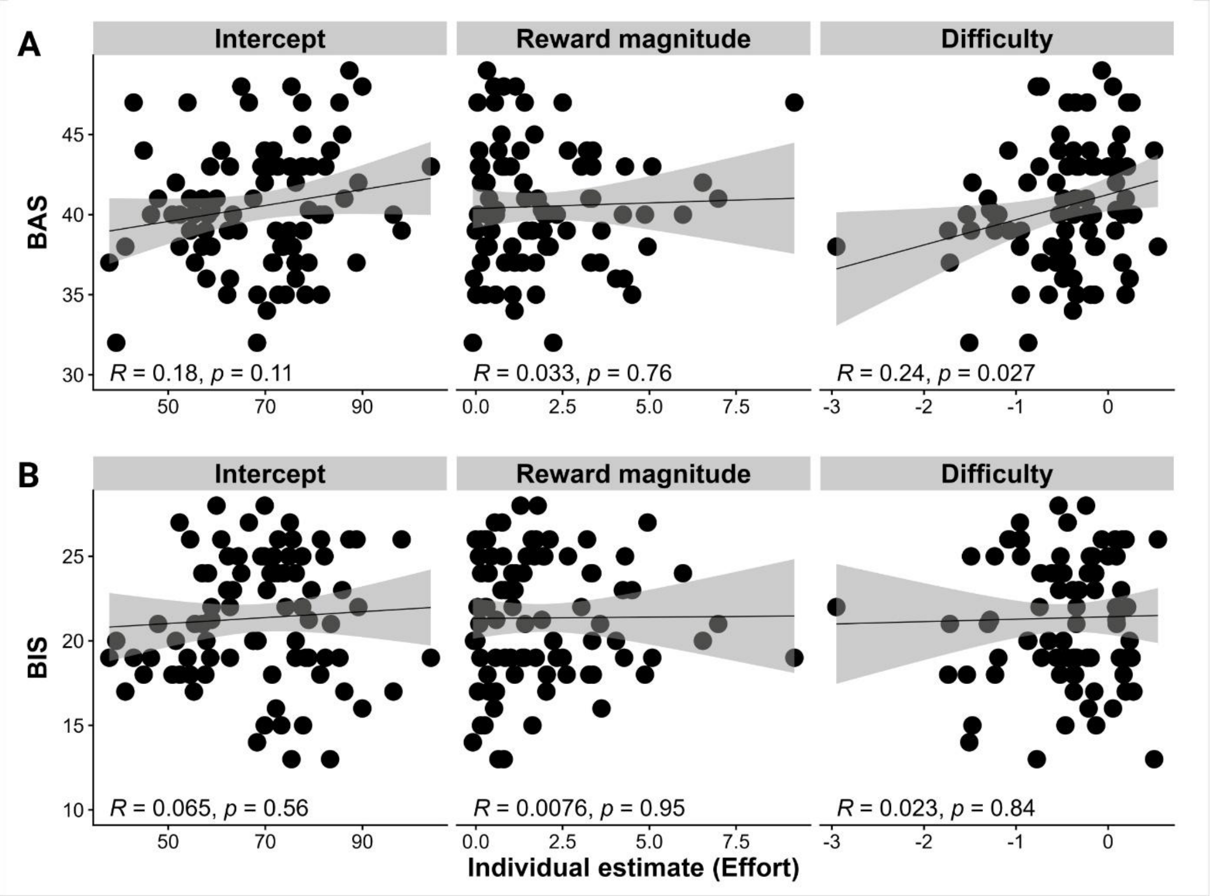


**Figure S6. Associations between behavioral effort indices and BIS/BAS scores.** A: Scatterplots of correlations between behavioral effort indices estimated from linear mixed-effects model of effort exertion (intercept, reward magnitude sensitivity, difficulty sensitivity) and Behavioral Activation System (BAS) scores. Pearson correlations showed a significant association with difficulty ($r = 0.24$, $p = .027$). B: Scatterplots of correlations between behavioral effort indices estimated from linear mixed-effects model of effort exertion (intercept, reward magnitude sensitivity, difficulty sensitivity) and Behavioral Inhibition System (BIS) scores. No significant association was observed.

### Supplementary Material H: Whole-brain NAcc functional connectivity results

**Table S3. Whole-brain NAcc seed-based functional connectivity results for all predictor models.**

| Predictor | Measure | Direction | Largest cluster region | | | | | |
|---|---|---|---|---|---|---|---|---|
| | | | $k_E$ | $t_{max}$ | $p_{FWE}$ | x | y | z |
| Effort exertion | Intercept | + | **141** | **4.65** | **.041*** | **46** | **12** | **24** |
| | | - | 18 | 3.81 | .988 | 4 | -26 | 48 |
| | Reward magnitude | + | 39 | 3.86 | .780 | 50 | -26 | -2 |
| | | - | 10 | 3.94 | .999 | -16 | 8 | 40 |
| | Difficulty | + | 53 | 4.23 | 0.565 | 30 | 38 | 40 |
| | | - | **155** | **5.15** | **.028*** | **-4** | **10** | **-24** |
| Wanting ratings | Intercept | + | 56 | 4.36 | .527 | 32 | 14 | -34 |
| | | - | 65 | 4.09 | .411 | 36 | 38 | 36 |
| | Reward magnitude | + | 19 | 4.07 | 0.984 | -20 | 19 | 4.07 |
| | | - | 2 | 3.36 | 1 | 40 | 2 | 3.36 |
| | Difficulty | + | 13 | 3.60 | .997 | 48 | 2 | -28 |
| | | - | 10 | 3.61 | .999 | 20 | -78 | -48 |
| Questionnaires | BIS | + | 87 | 4.92 | .208 | -8 | -2 | -18 |
| | | - | 31 | 4.11 | .901 | -30 | -28 | 6 |
| | BAS | + | 74 | 4.35 | .312 | 12 | 58 | 34 |
| | | - | 27 | 3.62 | .941 | -32 | -50 | 8 |
| | LARS | + | 28 | 4.29 | .924 | 28 | 26 | 18 |
| | | - | 17 | 3.60 | .991 | -52 | 12 | 20 |
| | SHAPS | + | 27 | 3.91 | .933 | 2 | 22 | -12 |
| | | - | 18 | 4.19 | .988 | -34 | -18 | 32 |

*Note:* $k_E$ = cluster size in voxels; $t_{max}$ = peak t-value; $p_{FWE}$ = cluster-level family-wise error (FWE) corrected *p*-value. Residual degrees of freedom were 76 for the effort and wanting models and 77 for the BIS/BAS and LARS-e/SHAPS models. Clusters are reported at $p < .001$ uncorrected. Asterisk (*) indicates surviving FWE correction at $p < .05$. MNI coordinates (x,y,z) reflect the peak voxel of the largest cluster. All models included age, sex, BMI, and mean framewise displacement (FD) as covariates of no interest.
